# Impact of Choline intake during pregnancy on maternal cognition and hippocampal gene signature in old age

**DOI:** 10.1101/2025.05.22.655437

**Authors:** Qian Yee Woo, Bernett TK Lee, Lee Wei Lim, Jingtao Zhang, Ayumu Tashiro, Pheck Khee Lau, Guillaume Thibault, Yulan Wang, Valerie C.L. Lin

## Abstract

Women are twice as likely to have Alzheimer’s disease (AD) than men and multiparity has been suggested to be a risk factor for dementia. The present study evaluated whether the lack of certain nutrients during pregnancy influences cognition while pregnant and in old age in mouse model. Non-targeted NMR analysis revealed significantly lower levels of numerous plasma nutrients and metabolites including choline and its derivatives on gestation day 7 compared to day 1. Novel object recognition and Morris Water Maze tests revealed impaired cognition in pregnant mice compared to nonpregnant controls. Choline deprivation worsened the cognitive impairment during pregnancy and choline supplementation alleviated it. Furthermore, choline availability during pregnancy affected cognition and general health in old age, with mice given a choline-deficient diet during pregnancy performed more poorly. RNA-Seq analysis indicates lasting effect of choline intake during pregnancy on hippocampal gene signatures in old age. Choline deprivation was associated with more upregulation of proinflammatory genes, whereas choline supplementation showed upregulation of neuroprotective genes such as *Prl*, *Gh*, and hemoglobin (*Hba* and *Hbb* subunits). Together, the study shows that choline supplementation benefits cognitive health in women during pregnancy and in old age.

## Introduction

Dementia is a neurodegenerative disorder characterized by the progressive loss of neurons and cognitive decline. According to the World Health Organization, 50 million people are currently living with dementia, and the number is predicted to reach 152 million by 2050. Alzheimer’s disease (AD) is the most common type of dementia, accounting for 60%–70% of all dementia cases. Notably, women have twice the risk of developing AD compared to men [1–3]. The reason for the higher proportion of women with AD is not clear. Because age is a major risk factor for AD and women generally live longer than men, they are more likely to reach the age of onset. However, other researchers postulate there are specific biological mechanisms underlying brain changes, progression and symptom manifestation that contribute to AD in women.

It is widely reported that women experience adverse cognitive changes during pregnancy, the so called ‘pregnancy brain’ [4, 5]. This phenomenon has been associated with structural changes in the brain of pregnant women. A prospective study revealed substantial reductions in the volume of grey matter postpartum in first-time mothers compared to nulliparous controls, and this reduction was still observed at the 2-year follow-up [6]. Studies have shown the grey matter reductions can be accumulative in consecutive pregnancies, with positive association between the number of pregnancies and the risk of dementia or AD [7–11]. Pregnancy is also associated with other neurological and behavioral changes, such as post-partum depression [12, 13]. It is therefore plausible that neurological changes during pregnancy have long-term implications. A better understanding of how pregnancy is associated with cognitive impairment and AD risk will shed light on novel strategies for disease prevention.

During pregnancy, the high demand for nutrients for fetal development can impact the supply of nutrients to the mother. Choline is particularly important for the brain, as it is a component of phosphatidylcholine (PC), the most abundant phospholipid in the brain that is required for structural integrity and function [14, 15]. Choline is also a precursor for acetylcholine, a neurotransmitter essential for learning and memory [16, 17]. Although studies have mainly focused the health consequences on offspring, they did show a significant decline in blood choline and PC in expectant mothers [18–21] Moreover, choline supplementation during pregnancy was shown to improve AD symptoms across multiple generations of offspring in an AD mouse model [22]. Similarity, studies have shown dietary choline has neuroprotective effects [23]. Although the importance of dietary choline on prenatal development and postnatal cognition has been well established, there has been no study on the impact of choline deficiency on mother’s cognitive health or the risk of neurodegenerative diseases in old age. In this study, we used Nuclear Magnetic Resonance (NMR) to profile changes in the plasma levels of metabolites during pregnancy in a mouse model, and found significant decreases of many nutrients including choline related metabolites. We also investigated the impact of choline availability on cognitive function during pregnancy and in old age. Lastly, we assessed whether multiparity and choline deficiency during pregnancy have a lasting effect on gene regulation in the hippocampus.

## Results

### Pregnancy is associated with significant decreases in the plasma levels of metabolites, particularly choline-related metabolites

To determine if the high nutrients demand of the growing fetuses affects maternal nutrients supply, we compared plasma levels of metabolites between gestation day 1 (GD1) and GD 7.5 in C57BL/6J mice by non-targeted NMR. We also determined whether the changes in the plasma levels of metabolites between the two stages are due to fetal demand by comparing the wild type mice (WT) and progesterone receptor (PgR) mutant mice in which fetal growth is retarded due to impaired uterine development [24]. The score plot shows that the data points between GD1 and GD 7.5 are more separated in the WT mice compared to that in PgR mutant mice on the X axis (Supplementary Fig. 1A). Quantification of signal intensity showed significant decreases of 16 plasma metabolites out of 25 from GD1 to GD7.5 in WT mice (Fig. 1A). In contrast, only 3 metabolites were significantly reduced in PgR mutant mice (Fig. 1B), which had stunted fetal growth compared to WT mice (insets in Fig. 1A and 1B). These decreases in the levels of metabolites during pregnancy were also clearly seen in the OPLS-DA coefficient plots from the NMR spectra, in which consistent decreases of choline, and its related metabolites: glycerophosphocholine (GPC), phosphatidylcholine (PC), and dimethylglycine (DMG) are marked out in WT mice (Supplementary Fig. 1C). In contrast, little change was observed in the PgR mutant (Supplementary Fig. 1D). Together, the NMR data indicate a broad decrease in plasma metabolite levels, specifically choline-related metabolites, during pregnancy due to the demands of the growing fetuses.

**Fig. 1.**
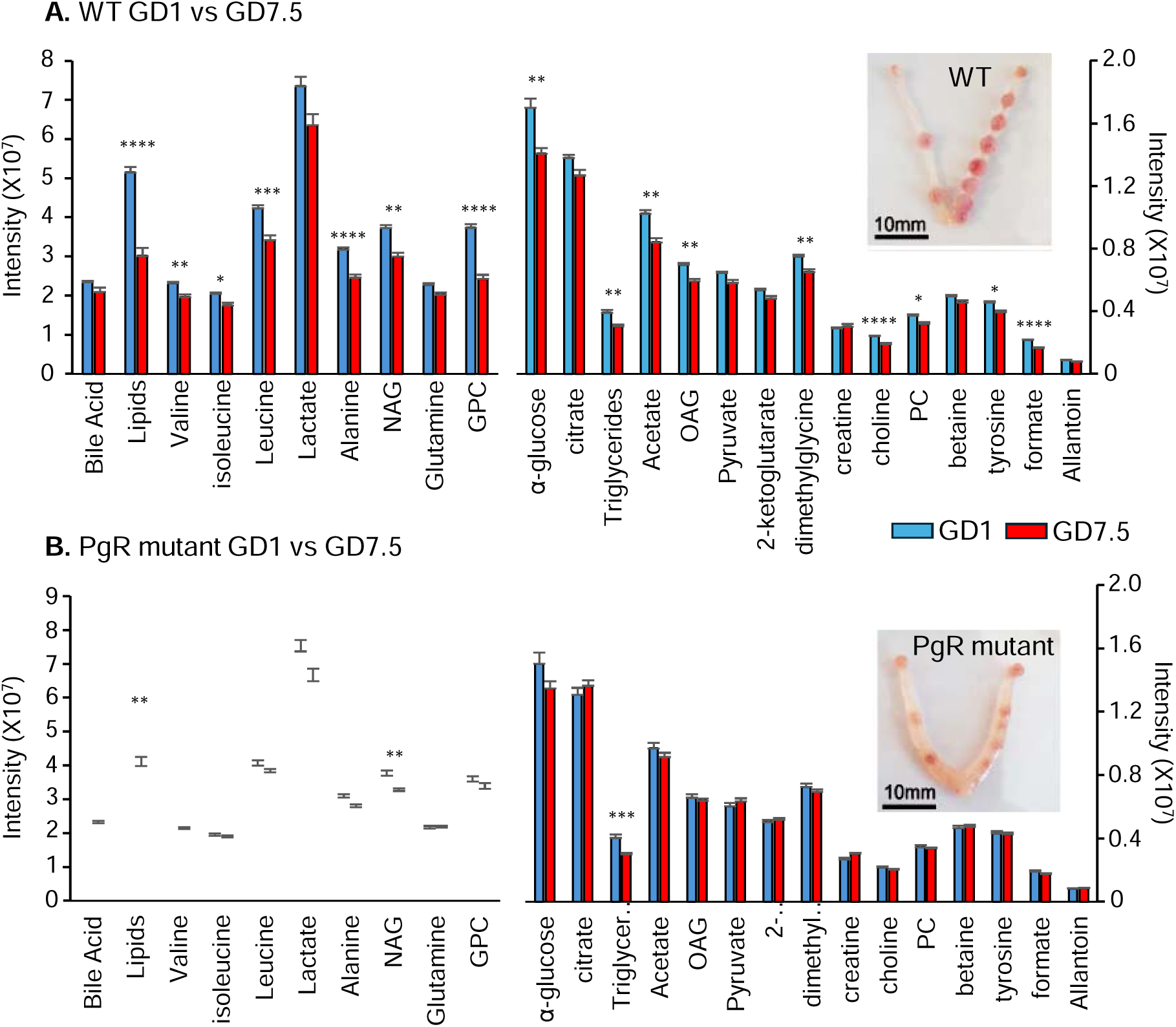
P**r**egnancy **is associated with significant decreases in the plasma levels of metabolites, particularly choline-related metabolites**. The bar chart shows NMR results of plasma metabolites on gestation day 1 (GD1, blue bar) and GD7.5 (red bar) for wild type (WT) **(A)** and Progesterone receptor (PgR) mutant mice **(B),** respectively. Differences in metabolite levels between GD1 and GD7.5 were analyzed using one-way ANOVA followed by Tukey’s post hoc test. Significantly more changes were observed in WT mice compared to PgR mutant mice. This is likely due to progressively increasing metabolic demands from growing fetuses in WT mice, which is not seen in PgR mutant mice due to stunted fetal development. The inset in the top-right corner of each bar chart shows fetal growth at GD7.5, highlighting the poor development of fetuses in PgR mutant mice compared to WT mice. All numerical data presented as mean ± SEM. Level of statistical significance: * p < 0.05, ** p < 0.01, *** p < 0.001, **** p < 0.0001.

The above findings indicate that there is a negative nutrients balance in pregnant mice on a standard laboratory diet. The reduced plasma levels of choline and related metabolites are particularly important in this context because people with lower plasma PC are at higher risk of developing AD and dementia [25–27]. While choline availability has been linked to offspring cognitive development [28–30], its effects on maternal cognitive health during pregnancy have not been studied. The present study tested the hypothesis that inadequate choline intake during pregnancy impairs mother’s memory and cognition during pregnancy and in old age. Four groups of mice were studied. These were Non-Pregnant control mice with a Normal diet (NP-N), Pregnant mice with a Normal diet (P-N), Pregnant mice with a Choline-Supplemented diet (P-CS) and Pregnant mice with an intermittent Choline- Deprivation (P-CD) (Fig. 2A). The P-CD mice were given normal diet on GD1, GD5 and GD9 to provide respite from choline deprivation. Pregnant groups were bred for four pregnancies (multiparity). Behaviour analyses were conducted during the second half of the first and fourth pregnancy, and at relatively old ages (12 and 15 months old) (Fig. 2B). The details of the test day during the pregnancy are shown in Fig. 2C. The hippocampus was collected for RNA-Seq analysis at 15 months old.

**Fig. 2.**
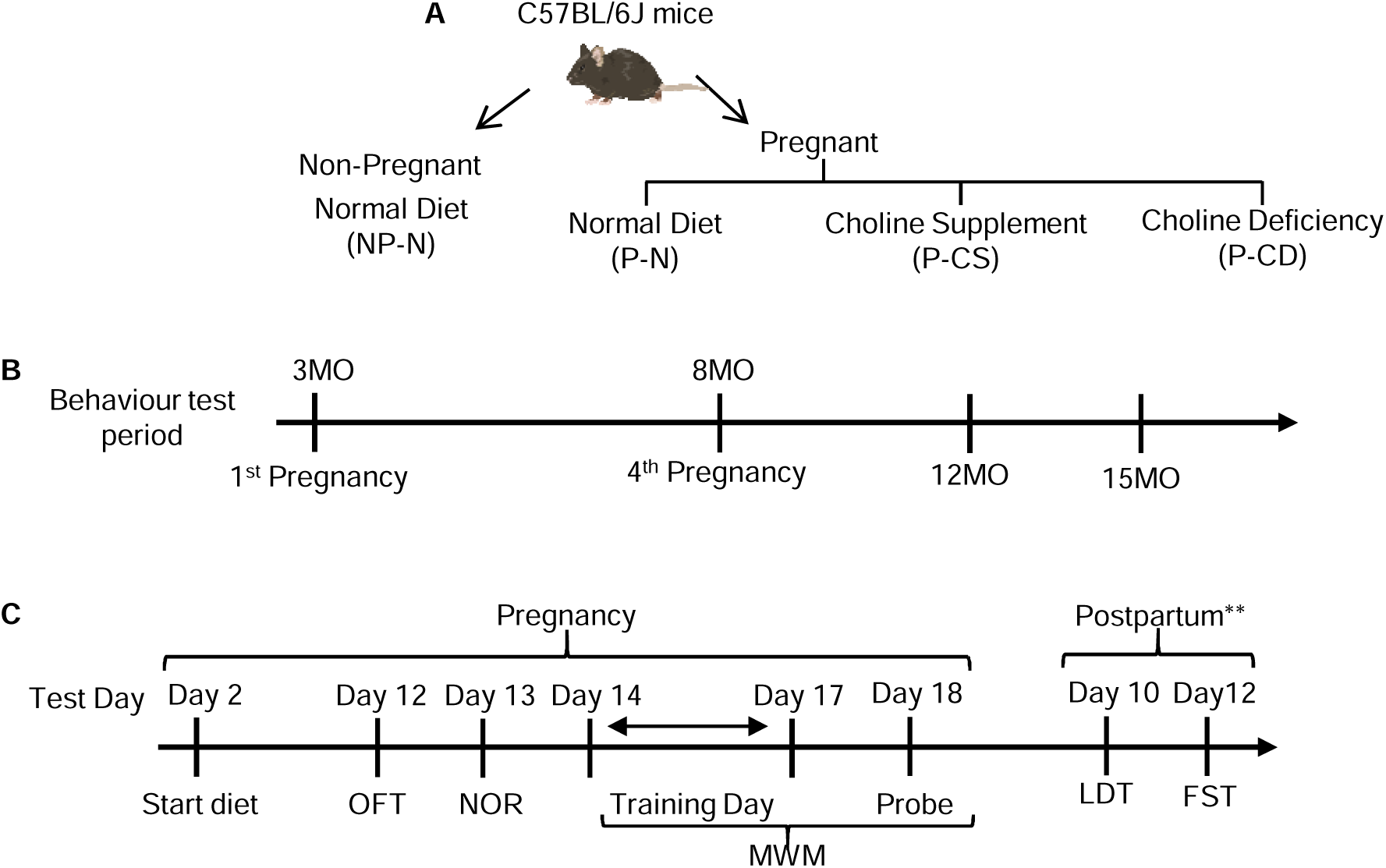
E**x**perimental **design for studying the effects of choline availability during pregnancy on memory (A)** C57BL/6J mice were randomly assigned to four groups: Non- Pregnant control with a Normal diet (NP-N), Pregnant mice with a Normal diet (P-N), Pregnant mice with a Choline-Supplemented diet (P-CS), and Pregnant mice with an intermittent Choline-Deprived diet (normal diet given on GD1, 5, and 9 during pregnancy) (P-CD). **(B)** Behavioral analyses were carried out during the second half of the first and fourth pregnancies, and at 12 and 15 months old. Open Field Test (OFT) and Novel Object Recognition (NOR) were conducted at 12 months, whereas NOR and Morris Water Maze (MWM) tests were carried out at 15 months. **(C)** During pregnancy, the specific diet was administered from GD2 to GD18. The OFT and NOR were conducted on GD12 and GD13, respectively. The MWM was conducted over five days, including four days of training and a probe session on the final day, from GD14 to GD18. ** Light/Dark Transition (LDT) test and Forced Swim Test (FST) were performed during the postpartum period after the fourth pregnancy, on postpartum day 10 and 12, respectively.

### Choline deprivation impairs aging-related performance in the open field test

The Open Field Test (OFT) was the first behaviour test, conducted on pregnancy day 12. The test measures anxiety, locomotor function, and exploratory behaviour. The OFT consists of a square arena marked with inner and outer zones. Rodents naturally spend more time in the outer zone, which is closer to the vertical wall [31]. The time spent in both zones and the total distance travelled were measured on day 12 of the first and fourth pregnancies, and at 12 months old. We performed the data analysis based on two categories: NP-N vs P-N mice and among pregnant mice (P-N vs P-CS vs P-CD).

There was no significant difference in the total distance travelled between NP-N and P-N groups and among the three pregnant groups in the first and fourth pregnancy, and at 12 months old (Fig. 3A), respectively. As expected, mice tended to spend more time in the outer zone than the inner zones (p < 0.0001), and there were no significant differences between groups, indicating anxiety levels were similar across all groups (Fig. 3B).

**Fig. 3.**
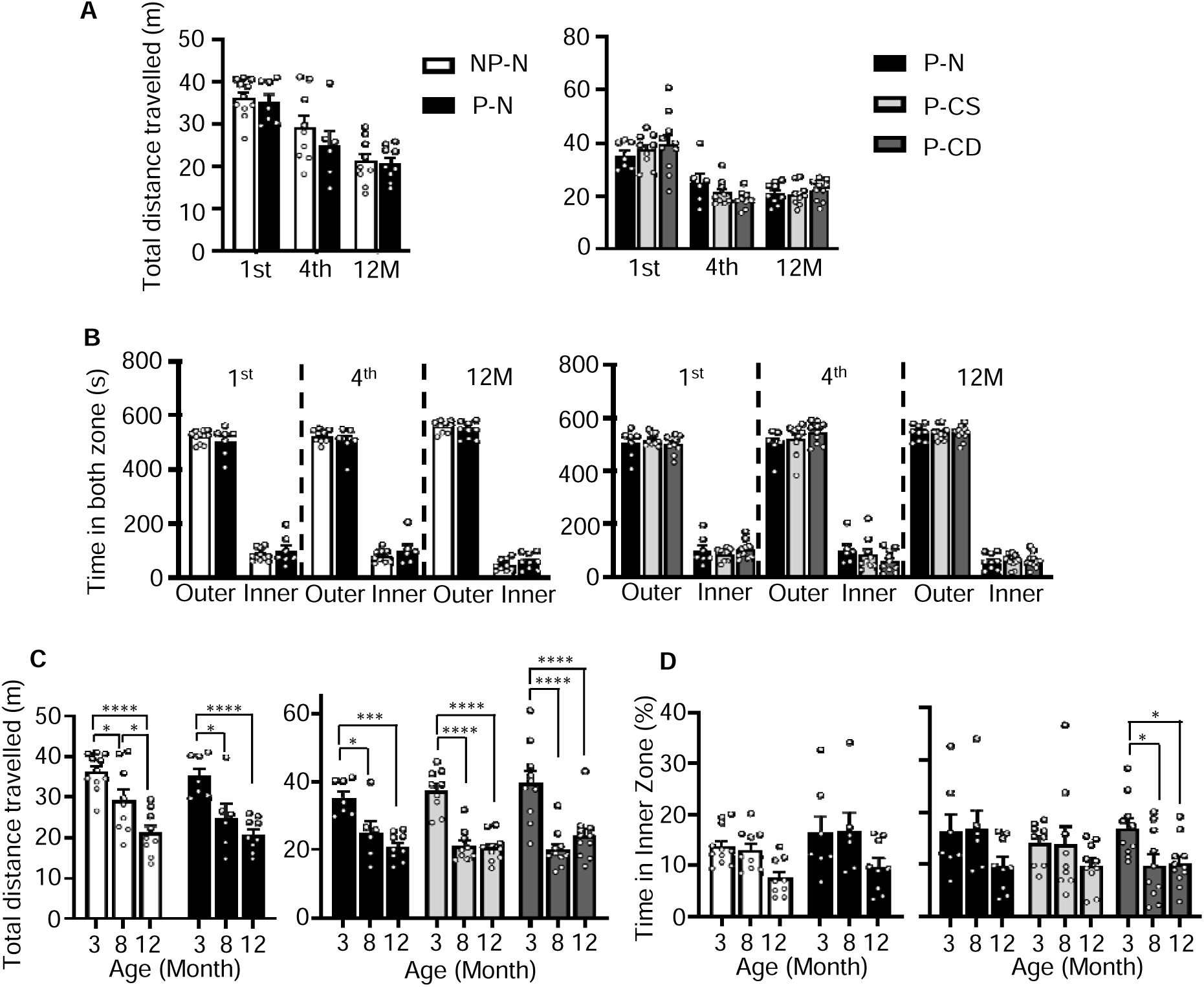
C**h**oline **deprivation impairs aging-related performance in the open field test. (A)** Bar chart of the total distance traveled by mice in the field box during the first and fourth pregnancies, and at 12 months of age. **(B)** Bar chart of the time spent in the outer and inner zones of the field box, which showed a significant difference between the time spent in both zones (p < 0.0001) across all groups. The results are the same for the first and fourth pregnancies, and at 12 months old, indicating no difference in the anxiety levels. **(C)** Total distance traveled was analyzed based on age in mice at 3 and 8 months old during the first and fourth pregnancies, respectively, as well as at 12 months of age. The results showed mice in all groups travelled significantly less as they aged. **(D)** Bar chart of the time spent in the inner zone based on age. P-CD mice at 8 and 12 months of age spent significantly shorter time in the inner zone compared to at 3 months of age. Choline deficiency appeared to impair mice’s exploratory behavior in the inner zone. All numerical data presented as mean ± SEM. Level of statistical significance: * p < 0.05, *** p < 0.001, **** p < 0.0001.

We also analysed the data based on the age, with the first pregnancy occurring at 3 months old and the fourth pregnancy at 8 months old. It was found that the total distance travelled in all groups decreased progressively as mice aged (p < 0.0001) (Fig. 3C). A significant age effect was found in the time spent in the inner zone (p = 0.0027). In particular, P-CD mice spent significantly less time in the inner zone at 8 and 12 months compared to mice at 3 months old (p = 0.0285 and p = 0.0337, respectively) (Fig. 3D). This indicates that choline deficiency caused impaired exploratory behaviour as mice aged.

### Choline deprivation impairs performance in the Novel Object Recognition test

The novel object recognition (NOR) test measures non-spatial memory in an open field. Rodents tend to explore a novel object rather than a familiar one. If their memory is intact and they can recognize the familiar object, they will spend more time exploring the novel object. Conversely, if they have cognitive deficits, there will be no difference in the time spent exploring the familiar and novel objects. The NOR test was carried out during the first and fourth pregnancies (GD13) and at 12 and 15 months old.

The percentage of time spent with the familiar and novel objects in each group of mice was analyzed. The NP-N mice group spent significantly more time with the novel object during the first and fourth pregnancies, as well as at 12 months of age (p < 0.0001) (Fig. 4A). However, at 15 months of age, the NP-N group no longer showed a significant preference for the novel object, suggesting cognitive decline in old age (Fig. 4A). The P-N mice spent significantly more time with the novel object in old age, both at 12 months (p = 0.008) and 15 months (p = 0.042); however, this result was not observed during pregnancy (Fig. 4B). This outcome may be due to low maternal choline levels during pregnancy, which could have affected cognitive performance. The P-CS mice spent significantly more time with the novel object during the first pregnancy and again at 15 months of age (1st: p < 0.0001; 15M: p = 0.0036) (Fig. 4C). Significantly more time spent with the novel object was also observed in P-CD mice (p = 0.0031) during first pregnancy. However, unlike the other groups, P-CD mice no longer showed a significant result for the novel object in subsequent time points (Fig. 4D). This may be due to the cumulative effects of choline deficiency into old age.

**Fig. 4.**
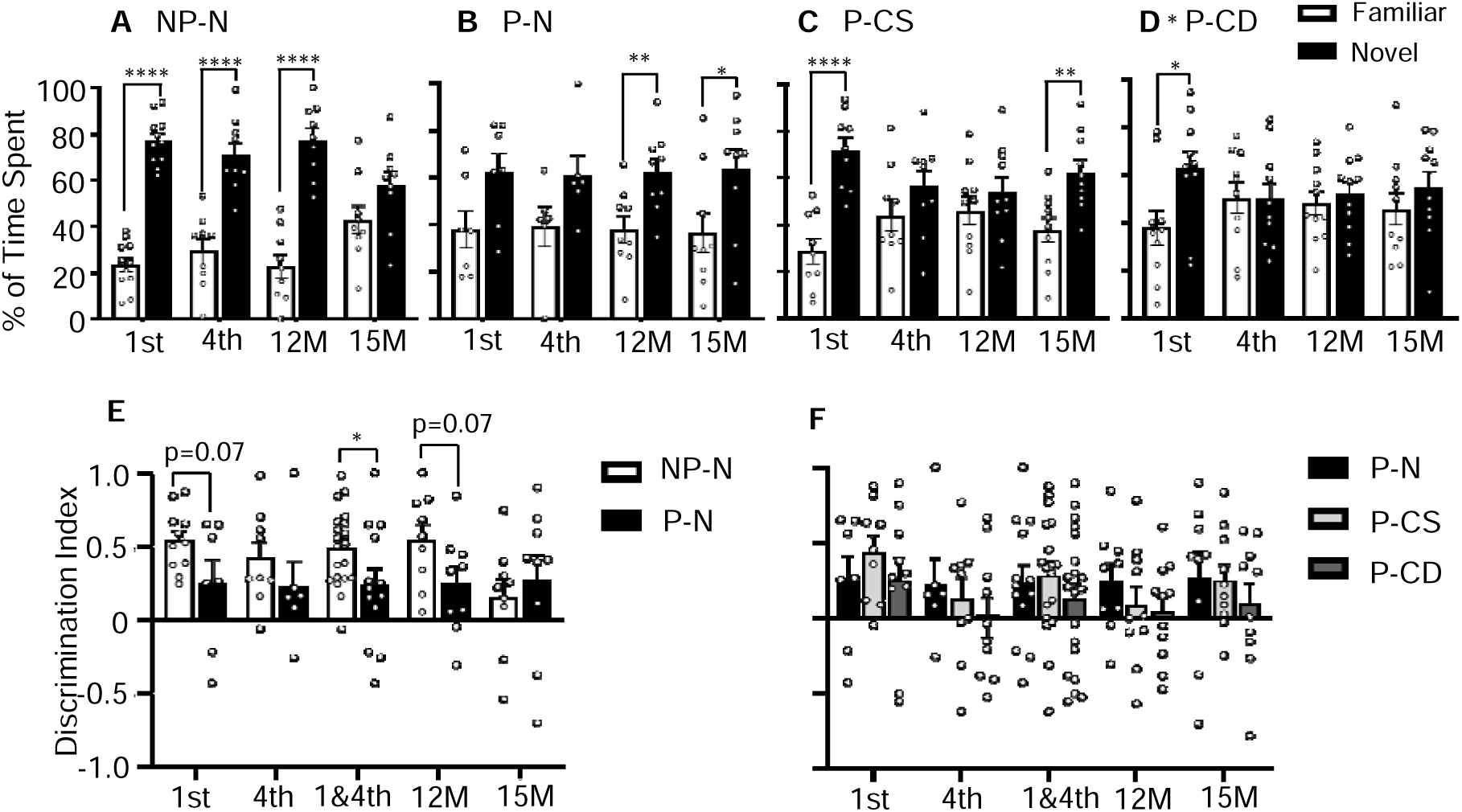
C**h**oline **deprivation impairs performance in the Novel Object Recognition test. (A, B, C, D)** The percentage of time spent on the familiar and novel objects during the first and fourth pregnancies, and at 12 and 15 months of age for the four different groups. Significant differences were observed in NP-N mice **(A)** (except at 15 months). P-N mice **(B)** only showed significant during their old age. P-CS mice **(C)** spent significant more time in novel object during first pregnancy and 15 months old. P-CD mice **(D)** mice failed to recognize the novel object except in first pregnancy only. **(E)** A bar chart of the discrimination index (novel object – familiar object / novel object + familiar object) showed a marginal difference between NP-N and P-N mice, except at 15 months of age, where no difference was observed. **(F)** A bar chart of discrimination index showed no significant difference between the pregnant groups. All numerical data presented as mean ± SEM. Level of statistical significance: * p < 0.05, ** p < 0.01, *** p < 0.001, **** p < 0.0001.

We also compared the discrimination index between NP-N and P-N, calculated as (Novel object – familiar object)/(Novel object + Familiar Object). NP-N mice appeared to perform better than the P-N mice during the first and fourth pregnancy, and at 12 months old (Fig. 4E) with marginal statistical significance (p=0.07 for 1^st^ pregnancy and 12 month; p=0.04 if data from the 1^st^ and 4^th^ pregnancies are combined). At 15 months old, NP-N mice no longer performed better in the NOR test (Fig. 4E). On the other hand, there was no difference in discrimination index among the pregnant groups at all the time points (Fig. 4F). Taken together, the NOR test results indicate that pregnancy is associated with cognitive deficits, and choline deprivation exacerbates this effect.

### Choline supplement during pregnancy improves memory in the Morris Water Maze test

Morris Water Maze (MWM) is commonly used to assess spatial learning and memory in rodents [32]. Mice were tested in MWM during the first and fourth pregnancies, and at 15 months old. Mice were administered four continuous training days, and a probe session on the fifth day.

The improvement in escape latency during the four training days indicates progressive learning. The P-CS mice performed the best during training days by taking a shorter time to find the platform than the previous day. The significant difference can be observed from training day 1 to subsequent three training days in the first and fourth pregnancy (p < 0.05) (Fig. 5A, 5B). Although NP-N mice did not show a significant reduction in escape latency over the four training days in the first pregnancy (Fig. 5A), they performed better in the fourth pregnancy, with significantly lower escape latency on day 3 and day 4 compared to day 1 (p = 0.0013, 0.0424) (Fig. 5B). The P-CD mice also showed lower escape latency on day 4 compared to day 1 in the fourth pregnancy (p = 0.0096). On the other hand, P-N mice did not show significantly different escape latency over the four training days.

**Fig. 5.**
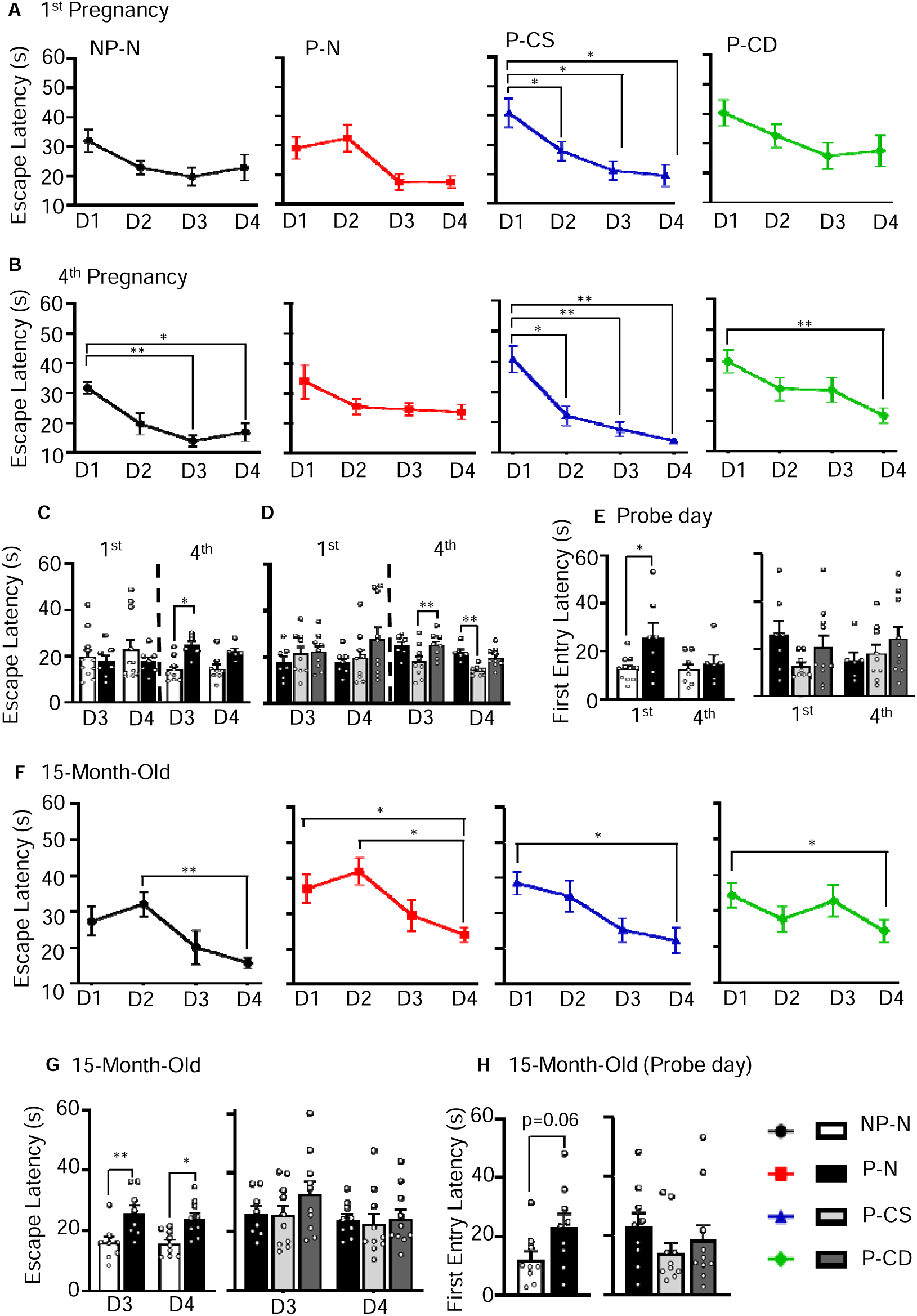
C**h**oline **supplement during pregnancy improves memory in the Morris Water Maze. (A, B)** Line graphs show the escape latency to find the hidden platform over four consecutive training days for the different groups during the first **(A)** and fourth **(B)** pregnancies. The significant differences in escape latency between training days reflect the learning progression of the mice throughout the training session. P-CS mice consistently showed a good performance, as indicated by significant differences between subsequent training days and the first training day. NP-N mice performed better during the fourth pregnancy compared to the first pregnancy. **(C)** Bar chart comparing escape latencies on training days 3 and 4 between NP-N and P-N mice. Significant difference was found on day 3 during fourth pregnancy only. **(D)** Bar chart comparing escape latencies on training day 3 and 4 between pregnant groups. Significant shorter latency was found in P-CS mice on both day 3 and 4 during fourth pregnancy. **(E)** The bar chart shows the latency to first entry to the platform on probe day. NP-N mice showed significant lower latency than P-N mice in first pregnancy. No difference was found between pregnant groups. However, there was a trend of decreasing latency to find the platform in P-CS mice during the first pregnancy. **(F)** The line graph shows escape latency during the training days at 15 months of age. **(G)** The bar chart shows escape latency on training days 3 and 4 at 15 months, which revealed significant different between NP-N and P-N mice on both training days. However, no differences found between pregnant groups. **(H)** Latency to first entry to the platform on probe day was marginally different between NP-N and P-N mice. Pregnant groups showed no significant differences. However, there was a trend of longer time to first entry in P-N and P-CD mice. All numerical data presented as mean ± SEM. Level of statistical significance: * p < 0.05, ** p < 0.01, *** p < 0.001, **** p < 0.0001.

Comparing escape latencies between NP-N and P-N mice on training days 3 and 4, no differences were observed during the first pregnancy. However, in the fourth pregnancy, NP- N mice exhibited significantly shorter escape latencies than P-N mice on day 3 (p = 0.012) (Fig. 5C). Although no difference was found during the first pregnancy, P-CS mice showed better performance in the fourth pregnancy, with a significant difference observed on training day 3 compared to P-CD mice (p = 0.005). A significant difference was also found on day 4, when P-CS mice were compared to both P-N (p = 0.0063) (Fig. 5D).

Swimming speed is one of the parameters used to assess motor impairments, which may potentially affect cognitive performance. No significant differences were found in swimming speed between groups during the first and fourth pregnancies (Supplementary Fig. 2A, 2B), although there was a trend suggesting that the mice tended to swim slower over the course of the training day. It is possible that the mice became more familiar with the task and experienced less stress, leading to slower but steadier swimming, as has been suggested [33].

The latency to first entry into the platform zone on the probe day for the first and fourth pregnancies is shown in Fig. 5E. NP-N mice showed significantly lower latency compared to P-N mice during the first pregnancy (p = 0.022). No significant differences were found in the latency between pregnant groups, although there was a trend of a better performance in finding the platform in the first pregnancy in P-CS mice compared to P-N and P-CD mice.

At 15 months old, the escape latency on day 4 was significantly lower compared to day 2 in NP-N mice (p = 0.0029) (Fig. 5F). On the other hand, P-N mice have significant lower escape latency on training day 4 compared to day 1 and 2 (p = 0.0188, 0.0104), respectively (Fig. 5F). P-CS and P-CD mice have significant escape latency on day 4 compared to day 1 (p = 0.0268, p = 0.0298). However, looking at the overall learning trend across the four training days, the P-CS mice showed a progressive reduction in escape latency, whereas the performance of the P-CD mice was not stable (Fig. 5F). NP-N mice outperformed P-N mice on both training day 3 (p = 0.0061) and 4 (p = 0.0144) at 15 months old (Fig. 5G). Although no significant differences were found on the third and fourth training days among the pregnant groups, there was a trend of shorter escape latency in the P-CS mice (Fig. 5G). Although a significantly slower swimming speed was observed in P-CD mice on training days 2 (Supplementary Fig. 2C), it was not associated with longer escape latency, as seen in the escape latency line graph for P-CD mice in Fig. 5F. Furthermore, the swimming speed of P-CD mice was not consistently slower than that of the other groups throughout the training days. Thus, we believe that the performance of P-CD mice during the training period was not affected by their motor function.

The latency to first enter the platform during probe day was marginal different (p=0.06) between NP-N and P-N mice (Fig. 5H). Similar results were observed among pregnant groups, with P-CS mice taking a shorter time for first entry to the platform than P-N and P-CD mice (Fig. 5H), although this did not reach statistical significance. Taken together, in MWM, the P-CS group performed the best across training days during pregnancy period, whereas both NP-N and P-CS groups performed better than P-N and P-CD mice on probe day, respectively. Although the learning trend during old age did not show significant variation among the groups across the training days, the NP-N and P-CS mice still demonstrated better performance, even though some of the results were not statistically significant.

### All pregnant groups exhibited postpartum anxiety in the light-dark test

We also tested the effects of pregnancy and choline availability on anxiety and depression levels during the postpartum period. The Light-Dark Test (LDT) and Forced Swim Test (FST) were conducted on postpartum days 10 and 12 of the fourth pregnancy, respectively.

The LDT test involves a box divided into a brightly lit, open zone and a dark, enclosed zone. Mice naturally prefer the dark compartment, with anxiety increasing with time spent in the dark zone. The cumulative duration spent in the light and dark zones is shown in Fig. 6A. In the test, NP-N mice spent almost similar time in the light and dark zone, while P-N mice spent significant longer time in dark zone (p < 0.0001). Regardless to the diet, all pregnant group spent significant more time in the dark zone (P-N: p = 0.001, P-CS: p = 0.0004, P-CD: p = 0.0069) (Fig. 6A). Fig. 6B showed the cumulative duration of mice spent in the light zone. NP-N mice spent significant longer duration in light zone compared to P-N mice, but no difference was found between the pregnant groups. The number of crossings between light and dark zone was no difference between NP-N and P-N, or between pregnant groups (Fig. 6C).

**Fig. 6.**
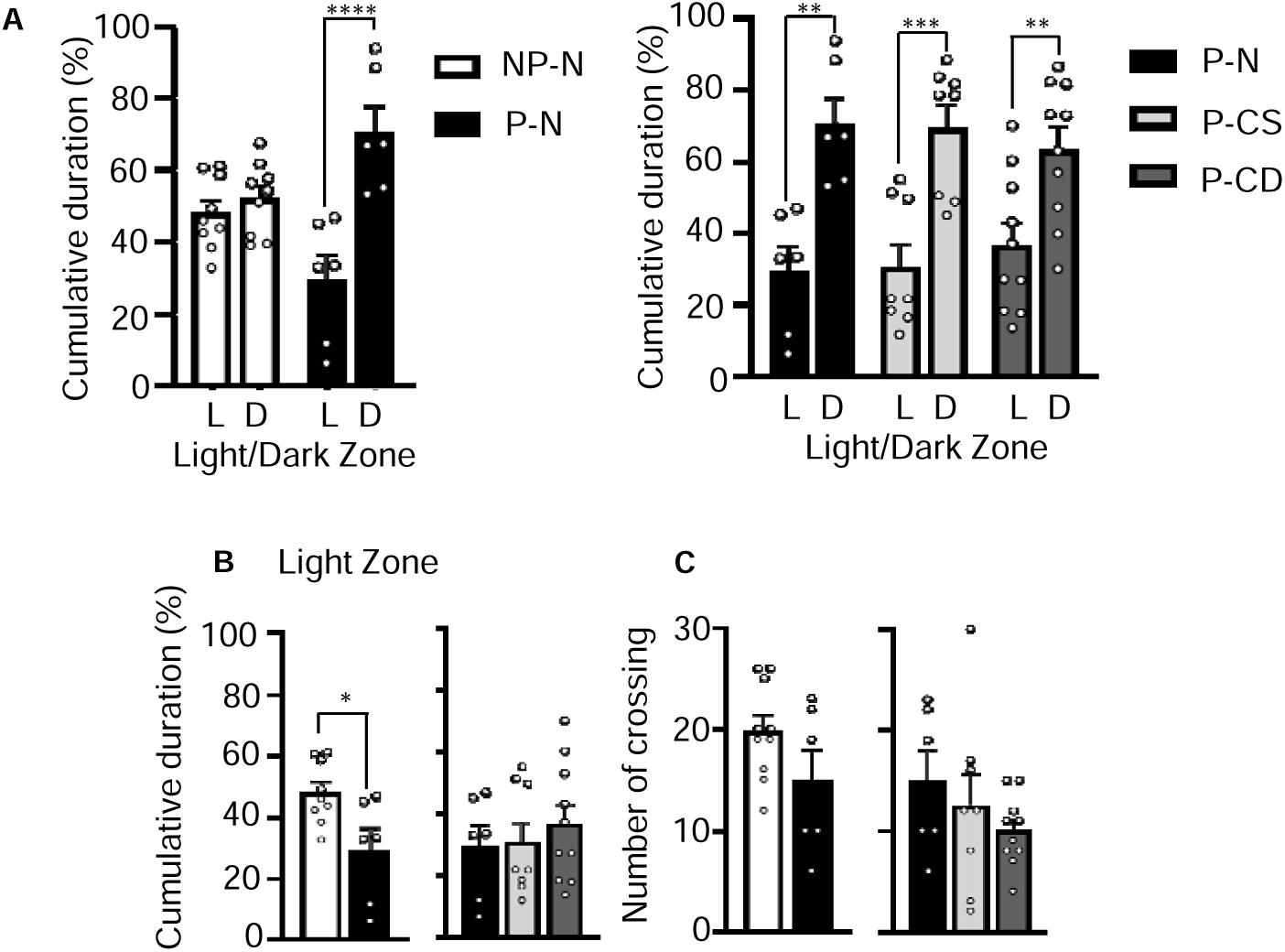
A**l**l **pregnant groups exhibit postpartum anxiety in the light-dark test. (A)** Cumulative duration spent in the light and dark zones during the light-dark test. Significant differences were found in all pregnant groups, indicating that the pregnant mice experience anxiety and tend to hide in the darker area. **(B)** The bar chart showed the cumulative duration of mice spent in light zone. Significant longer time spent was found in NP-N mice compared to P-N mice. No significant difference found between pregnant groups. **(C)** The bar chart shows the number of crossings between the light and dark zones during the light-dark test among the groups. Data are presented as mean ± SEM. The degree of statistical significance is indicated by asterisks (* p < 0.05, ** p < 0.01, *** p < 0.001, **** p < 0.0001).

The forced swim test did not reveal any significant differences in immobility duration or the frequency of immobility (Supplementary Fig. 3A, 3B). Hence the degree of depression was similar among groups.

### Choline availability during pregnancy influences body weight and skin condition during pregnancy and in old age

We determined whether choline deprivation or supplementation affects metabolism and hence body weight on day 1, 10 and 17 of pregnancy. During the first, second and fourth pregnancy, P-CD mice was significantly heavier than the P-N mice with normal diet on day 10 and 17 of pregnancy with the most striking effect on the second pregnancy (day 10: p = 0.0004, day 17: p < 0.0001) (Fig. 7A, 7B and 7C). P-CS mice also significantly increased body weight compared with P-N mice on the second pregnancy (day 10: p = 0.0205, day 17: p < 0.0001) and fourth pregnancy (day 17: p = 0.0388) (Fig. 7B, 7C), although the difference on day 10 of fourth pregnancy was not statistically significant (Fig. 7C).

**Fig. 7.**
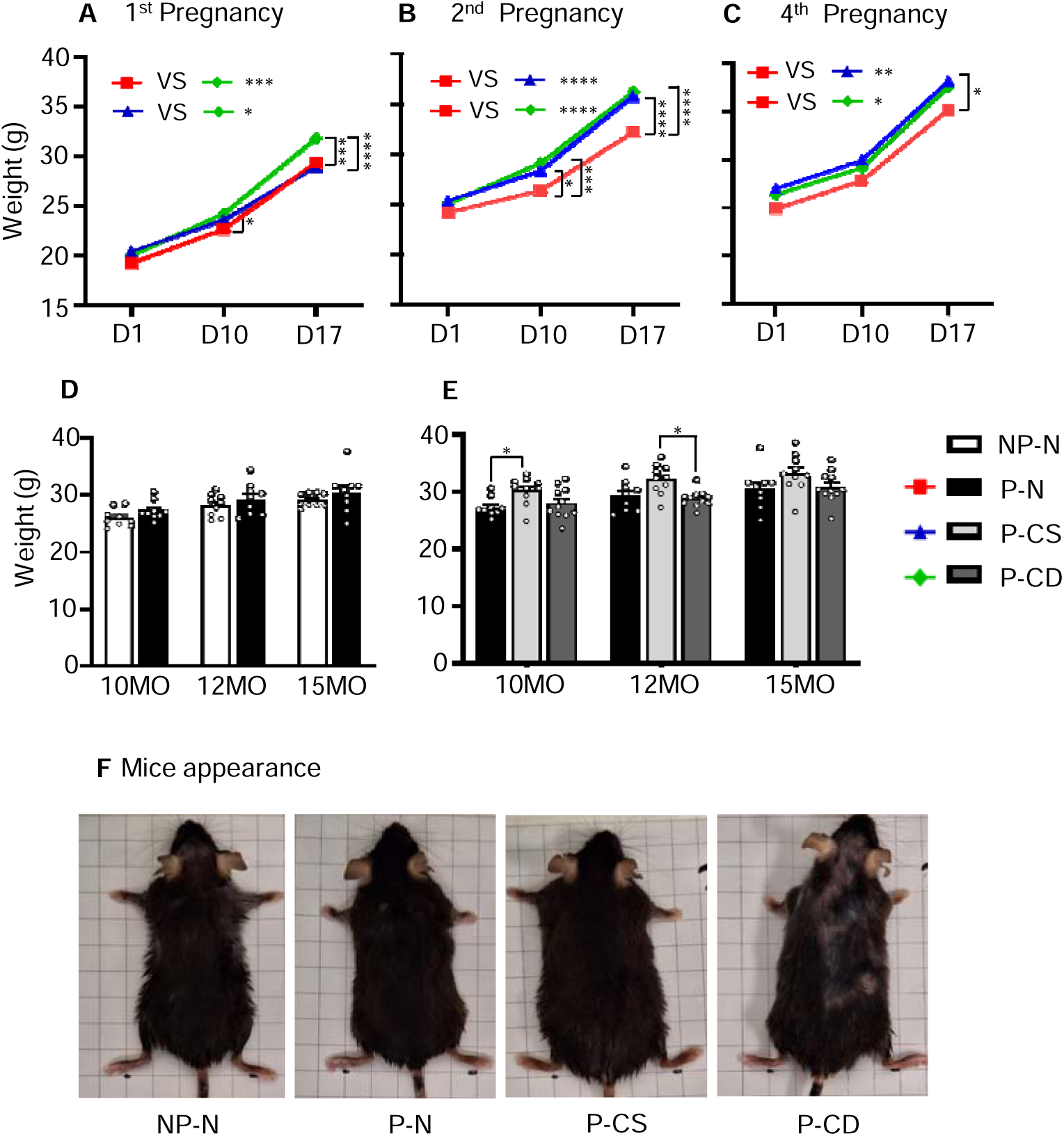
C**h**oline **availability during pregnancy influences body weight and skin condition during pregnancy and at old age. (A, B, C)** Mice weight on day 1, 10, and 17 during the first **(A)**, second **(B)**, and fourth **(C)** pregnancies. **(D)** The weights of NP-N and P-N mice during old age were not significantly different. **(E)** The weights of all pregnant groups during old age. Significantly higher weight was found in P-CS mice at 10 and 12 months of age compared to P-N and P-CD respectively, but no difference was observed at 15 months of age. **(F)** Pictures were taken at 15 months of age, just before sacrifice. P-CD mice had a poorer skin condition compared to the other groups. Data are presented as mean ± SEM. The degree of statistical significance is indicated by asterisks (* p < 0.05, ** p < 0.01, *** p < 0.001, **** p < 0.0001).

We also analysed mice weight after the fourth (last) pregnancy. At 10 and 12 and 15 months old, P-N mice had similar weight to NP-N mice (Fig. 7D). Among the three pregnant groups, P-CS mice were generally heavier, with significant differences between P-CS and P-N at 10 months old (p = 0.0478) and between P-CS and P-CD at 12 months old (p=0.0122) (Fig. 7E).

Lack of choline during the pregnancy negatively impact skin health. In general, P-CS mice looked healthier with shiny skin hairs. Choline deprivation during pregnancy resulted in a skin condition with hair loss resembling that of alopecia. This happened in every pregnancy and the symptoms persisted in old age (Fig.7F, Supplementary Fig. 4).

### Pregnancy and choline supply has lasting effect on hippocampal gene expression

We have shown that pregnancy negatively influences cognitive behavior, whereas choline supplement partially rescued these negative effects. This suggests that choline availability during pregnancy has long-term effects on the body. We performed RNA-Seq analysis on 15-month-old mice to evaluate whether choline availability during pregnancy has long-term effect on gene signatures in the hippocampus, the memory hub of the brain. In addition to the four experimental groups, we also included 8-week-old young mice in the gene expression analysis to compare ‘young’ and ‘old’ gene expression signatures.

#### Aged mice show dysregulated of genes related to neuroinflammation, growth, and memory compared to young mice, whereas pregnancies partially reversed the dysregulation

As expected, there were significant differences in the gene expression between young mice and old nulliparous mice (8 weeks vs 15 months). There are 259 differential expressed genes between young and old nulliparous mice (p < 0.01), of which 112 were upregulated and 147 were downregulated. The differentially expressed genes between the young and old multiparous mice showed 30% overlap with differential genes between nulliparous young and old mice (Fig. 8A), suggesting that multiparity influences hippocampal gene expression.

**Fig. 8.**
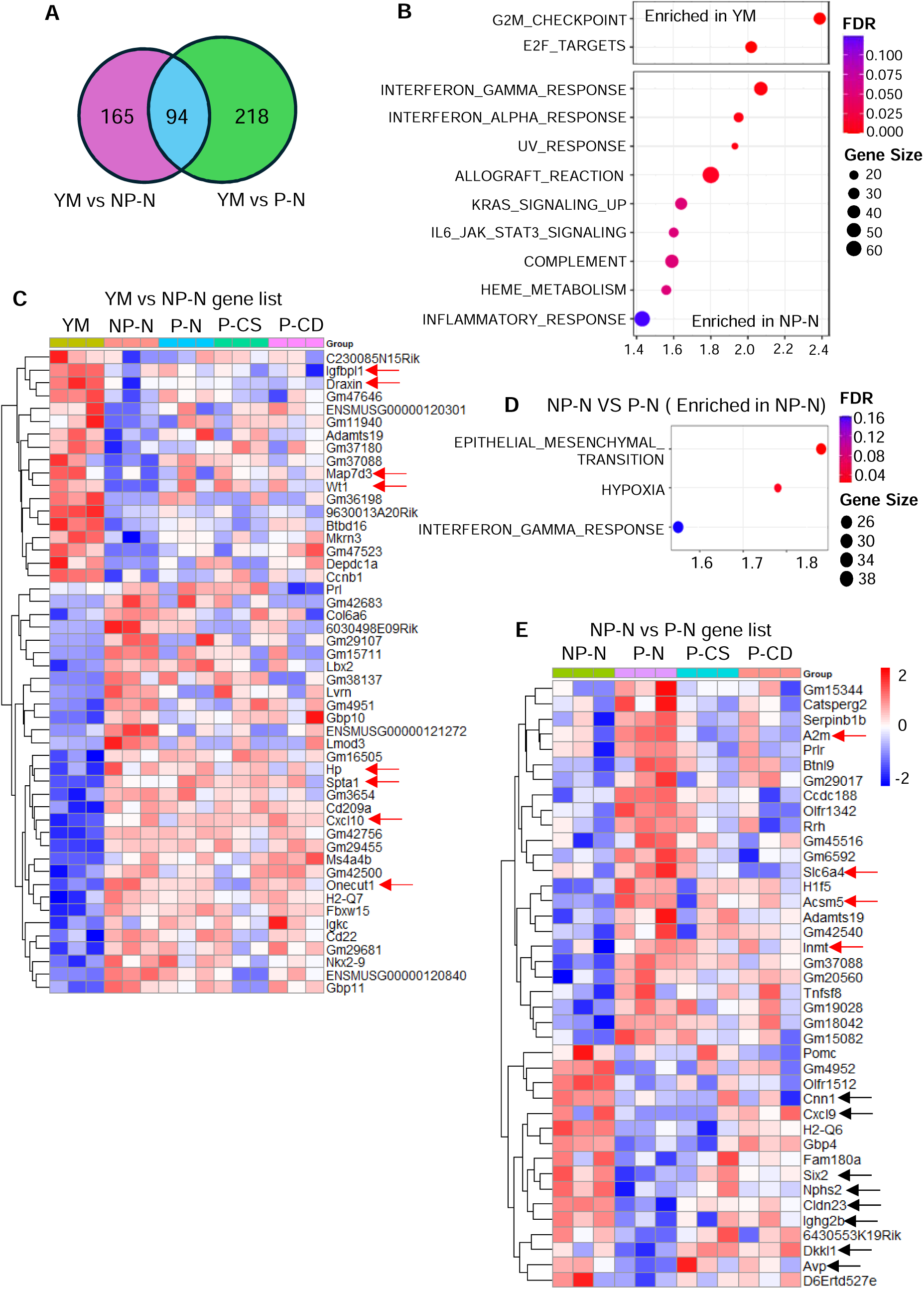
RNA-Seq analysis shows differential gene expression patterns in the hippocampus between the young (YM) and old nulliparous mice (NP-N) and old multiparous (P-N). **(A)** The Venn diagram shows the number of overlapping significant genes (n = 94) in both NP-N (old and nulliparous) and P-N (old and multiparous) compared to YM (8 weeks old young mice). **(B)** Heatmaps were plotted based on the list of significant genes (p < 0.01) for YM compared to NP-N (n = 50). **(C)** Heatmap of the significant genes list from the NP-N and P-N comparison (n = 40). **(D)** GSEA analysis reveals three enriched hallmarks in NP-N (p < 0.05, FDR < 0.25), but no enrichment was found in P-N. **(E)** The significant genes list from the NP-N and P-N comparison (n = 40), which are more consistent among the replicates, is summarized in the heatmap. The red and black arrows show notable genes to take note.

Gene Set Enrichment Analysis (GSEA) from the Molecular Signature Database (MSigDB) [34–36] showed that the hippocampus of young mice is enriched with proliferative hallmarks G2M_checkpoint and E2F-targets compared to old mice (Fig. 8b). In contrast, 6 of the top 10 enriched hallmarks in old mice are related to immune reactions including Interferon_Gamma_Response (proinflammatory) and Interferon_Alpha_Response (anti- inflammatory), Allograft_Reaction and IL6_JAK_STAT3_signaling.

Of the 259 differentially expressed genes between young and old nulliparous mice, we plotted a heatmap of 50 genes that showed the most consistent changes among the triplicates (Fig. 8C). Downregulated genes in old nulliparous mice were generally also downregulated in multiparous groups, albeit to a lesser extent. Among these downregulated genes, *Igfbpl1*, *Draxin*, *Map7d3*, and *Wt1*, which are indicated by red arrows, are known to have neural functions. *Igfbpl1* (Insulin Like Growth Factor Binding Protein Like 1) plays a crucial role in microglia homeostasis, is involved in resolving lipopolysaccharide-induced neuroinflammation and tauopathy [37], and regulates axonal growth [38]. *Draxin* is an axon guidance protein essential for brain development, including forming neural circuits and guiding axonal growth and migration [39]. *Map7d3* (MAP7 Domain Containing 3) encodes a microtubule-associated protein primarily expressed in the brain, and its mutation is linked to intellectual disability and autistic traits[40]. *Wt1* (Wilms’ tumor 1) gene is involved in neural progenitor cell proliferation and neuronal differentiation [41], and also plays an important role in memory flexibility by limiting memory strength or promoting memory weakening [42]. Together, downregulation of these genes in old mice results in increased neuroinflammation and reduced neural growth and progenitor activity. Interestingly, these genes were less downregulated in old multiparous mice compared to nulliparous mice, suggesting that pregnancies offer protection in the brain as mice age.

Similarly, genes upregulated in old nulliparous mice were also generally upregulated in multiparous mice groups with some exceptions (Fig. 8C). Notable genes that were consistently upregulated in old mice compared to young mice include *Hp*, *Spta1*, *Cxcl10*, and *Onecut1*, which are indicated by black arrows and are involved in various aspects of neural function and neuroinflammation. Haptoglobin (*Hp*) is a hemoglobin-binding protein that alleviates iron-related oxidative damage, whereas its deletion in old mice led to greater brain lesions in a traumatic brain injury model [43, 44]. *Spta1* (Spectrin Alpha, Erythrocytic 1) gene encodes a cytoskeletal protein, and is crucial for maintaining the structural integrity of cell membranes, particularly in neurons [45]. The proinflammatory cytokine *Cxcl10* (CXC Motif Chemokine Ligand 10) plays a part in the CXCL10/CXCR3 axis, which involved in the induction of cerebral hyperexcitability in response to an inflammatory trigger [46–48]. *Onecut1* is an atypical homeodomain transcription factor required for neuronal characterization and horizontal cell determination [49] . *Onecut1*-null mice show defective synaptogenesis, and age-dependent degeneration of photoreceptors [50]. Taken together, upregulation of *Hp*, *Spta1*, *Cxcl10* and *Onecut1* in old mice is likely required to maintain neural integrity and perhaps memory/wisdom.

#### Differential gene expression between age matched nulliparous and multiparous old mice

Comparison of nulliparous (NP-N) and multiparous mice (P-N) data showed 199 differentially expressed genes in NP-N (p < 0.01), with 94 upregulated and 105 downregulated genes. GSEA analysis showed that Epithelial Mesenchymal Transition, Hypoxia, and Interferon Gamma response were the most significant hallmarks enriched in nulliparous mice (NP-N) (FDR < 0.25) (Fig. 8D). Upregulation of these processes means more stress, tissue remodelling, and inflammation in the hippocampus of nulliparous mice compared to multiparous mice. Gene set enrichment analysis did not yield significantly enriched hallmarks in the multiparous group, suggesting that the differentially regulated genes in this comparison are of diverse biological pathways instead of being enrich in specific hallmarks.

The heatmap in Fig. 8E shows 40 genes with the most consistent changes between nulliparous and multiparous groups. Notable upregulated genes in multiparous mice (red arrow) include *A2m*, *Slc6a4*, *Acsm5*, and *Inmt*. The *A2m* (Alpha-2-Macroglobulin) protein has been implicated in Alzheimer’s disease (AD) due to its ability to mediate the clearance and degradation of amyloid-beta [51], although its involvement is still controversial [52, 53]. The *Slc6a4* gene encodes a protein that transports serotonin from the synaptic cleft back into the presynaptic neuron. Polymorphisms in *Slc6a4* have been linked to differences in personality traits, and vulnerability to stress-induced symptoms [54, 55]. Studies on *Slc6a4* (5-Htt) knockout mice showed *Slc6a4* regulates neuronal morphology and plasticity in limbic brain areas in a repeated social defeat stress model [56]. The *Acsm5* gene, encodes Acyl-CoA Synthetase Medium Chain Family Member 5, is involved in fatty acid metabolism in the brain and inhibits ligamentum flavum hypertrophy by regulating lipid accumulation [57]. *Inmt* (Indolethylamine N-Methyltransferase) plays a role in the synthesis of N,N-Dimethyltryptamine (DMT), a psychedelic compound found in the mammalian brain [58]. Its deletion did not result in overt behavioral or physiological abnormalities in rats [59]. The upregulation of these genes in multiparous mice suggests that pregnancies offer some benefits to the function of hippocampus.

Notable upregulated genes in nulliparous mice versus multiparous mice (black arrow) included Cnn1, *Cxcl9*, *Six2*, *Nphs2*, *Cldn23*, *Ighg2b*, *Dkkl1*, and *Avp*, which are all involved in various neural functions (Fig. 8E). For example, the *Cxcl9* (CXC Motif Chemokine Ligand 9) is involved in recruiting immune T cells into the brain, particularly during the early stages of reactivation of chronic cerebral infection [60]. *Six2* (Six Homeobox 2) is a transcription factor that plays a role in regulating craniofacial development and frontonasal morphogenesis[61]. It also mediates the protective effect of Glial cell line-derived neurotrophic factor (GDNF) on dopaminergic neurons. *Six2* gene silencing led to increased apoptosis of damaged dopaminergic neurons [62]. Thus, higher levels of *Six2* in nulliparous mice would be beneficial.

#### Choline supplement is associated with upregulation of Hba, Hbb, Gh and Prl in hippocampus

There were significant differences in hippocampal gene expression between P-N and P-CS mice (Fig. 9A), and between P-N and P-CD mice (Fig. 9B). Intriguingly, choline-deprived and choline-supplemented mice had similar genes expression pattens as seen by similar heatmaps when compared to control mice on a normal diet. On the other hand, comparisons between P-CS and P-CD mice also showed significant expression differences in many genes (Fig. 9C). These findings suggest that choline supplementation or choline deprivation affects the expression of distinct sets of genes.

**Fig. 9.**
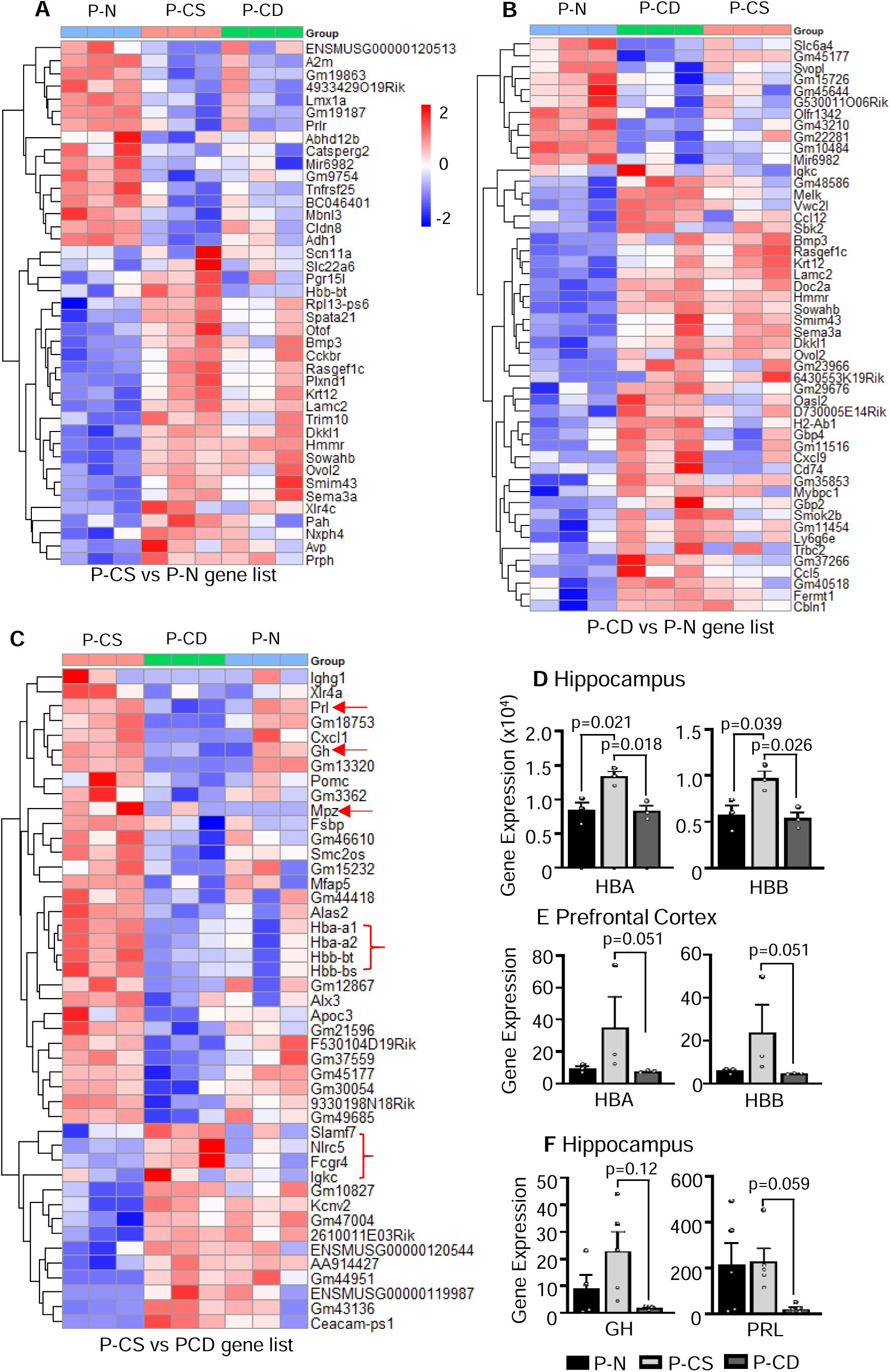
C**h**oline **availability during pregnancy is associated with altered hippocampal gene expression. (A, B)** Heatmaps of the significant gene lists (p < 0.01) for P-CS compared to P-N mice (n = 41) **(A)**, and P-CD compared to P-N mice (n = 50) **(B)**, respectively. **(C)** Heatmap of the comparison between P-CS and P-CD mice (n = 45) with p < 0.01 as the cut- off. The red arrows and right curly brackets show notable genes to take note. **(D)** Validation of RNA-Seq data by qPCR. Comparison between the pregnant groups revealed P-CS mice had the highest levels of *Hba* and *Hbb* in hippocampus compared to P-N and P-CD mice. **(E)** qPCR analysis on *Hba* and *Hbb* levels in prefrontal cortex. P-CS mice had marginally significant higher compared to the P-N and P-CD mice. **(F)** Validation of RNA-Seq data by qPCR. Comparison between the pregnant groups showed lower *Gh* and *Prl* levels in P-CD mice. All numeric data presented as mean ± SEM.

It is noteworthy that these mice were saline-perfused before brain collection, yet the expression of four haemoglobin genes *Hba-a1*, *Hb-a2*, *Hbb-bt*, *Hbb-bs* were significantly higher in the hippocampus of P-CS mice compared P-N and P-CD mice (Fig 9C). Haemoglobin genes upregulation in the P-CS mice hippocampus was validated by qPCR (Fig. 9D). Interestingly, *Hba* and *Hbb* expression were also higher in the prefrontal cortex of P-CS mice than that in P-N and P-CD mice (Fig. 9E). Neuronal Hbs were reported in neurons of several brain regions including the hippocampus and cortex and play a role in oxygen storage and transport [63–66]. Neuronal *Hba* and *Hbb* showed strong mitochondrial localization [67], presumably involved in energy metabolism. Several studies suggested that Hbs are involve in neurodegeneration and AD (see review in [68]). For example, lower HBA or HBB protein levels were detected in neurons with granular or punctuate hyperphosphorylated tau deposits, in AD neurons with tangles in the hippocampus and frontal cortex [69, 70]. Another study reported strong downregulation of *Hba* and *Hbb* mRNA in brains of AD patients [71]. The higher expression of Hbs in the P-CS hippocampus and prefrontal cortex suggest beneficial effects on choline supplement on brain health.

The expression of *Gh* (Growth hormone) and *Prl* (Prolactin) are also significantly higher in P-CS mice compared to P-CD mice (Fig. 9C). This was also validated in hippocampus by qPCR with marginal significance (Fig. 9F). *Gh* and *Prl* share significant sequence homology, structural similarities and overlapping functions [72]. Studies suggested that *Gh* upregulates Hbs’ concentration in response to oxygen or metabolic demands via IGF- 1[73]. This is supported by the evidence that Gh replacement in a Gh-deficient rat model enhanced the expression of hemoglobin (*Hba* and *Hbb*) in the hippocampus [74]. Consistent with this understanding, the upregulation of *Gh* is associated with upregulation of all Hbs in P-CS mice compared to P-CD mice. *Gh* is also known to be involved in the proliferation of neural progenitors and increased level of *Gh* was found to enhance the density of dendritic spines in hippocampus and better performance in behaviour tests [75]. Additionally, the age-related reduction in Gh secretion is considered to contribute to the neurogenic decline in the elderly and administration of Gh replacement has been shown to be effective in improving memory and learning activities [76, 77]. Similarly, Prl also plays a significant neuroprotective role in the hippocampus. It has been shown to enhance hippocampal synaptic plasticity, improve spatial learning, and exhibit neuroprotective effects against excitotoxicity [78–84]. Taken together, the upregulation of *Gh* and *Prl* in the hippocampus of P-CS mice is favourable for neuronal health.

*Nlrc5*, *Slamf7*, *Fcgr4* and *Igkc* genes were downregulated in the hippocampus of P-CS mice compared to P-CD mice (Fig. 9C). These genes are important for immune function. It was reported that *Nlrc5* (NLR Family CARD Domain Containing 5) promotes neuroinflammation and inhibits neuronal survival in conditions such as multiple sclerosis and Parkinson’s disease [85, 86]. Although direct evidence linking *Slamf7*, *Fcgr4*, and *Igkc* to neurodegenerative diseases is limited, their roles in immune modulation present plausible pathways for their involvement in CNS disorders [87–89].

#### Enrichment of pro-inflammatory gene sets in P-CD versus P-N mice hippocampus

Comparison between P-CD mice and P-N mice in GSEA analysis showed significant enrichment of immune function related hallmarks including TNFA_Signaling_Via_NFKB, Interferon_Gamma_Response, Interferon_Alpha_Response, Inflammatory_Response, and Allograft_Reaction (Fig. 10A). Intriguingly, the P-CS hippocampus is also enriched with these hallmarks compared with P-N. However, 24 genes were significantly (p < 0.05) upregulated in P-CD vs P-N mice from these five enriched hallmarks (Fig. 10B), as opposed to 7 genes in P-CS vs P-N mice (Fig. 10C), indicating a greater activation of immune response in P-CD than P-CS hippocampi. Since the brain’s immune system is commonly activated in response to injury or neurodegeneration, the upregulation of genes related to the immune response in P-CD mice suggests proinflammatory effects of choline deprivation during pregnancy. The upregulation of a few proinflammatory genes, such as *Il1b*, *Il12a*, and *Cd28*, in P-CS mice adds an additional layer of complexity to the role of these genes in neural biology.

**Fig. 10.**
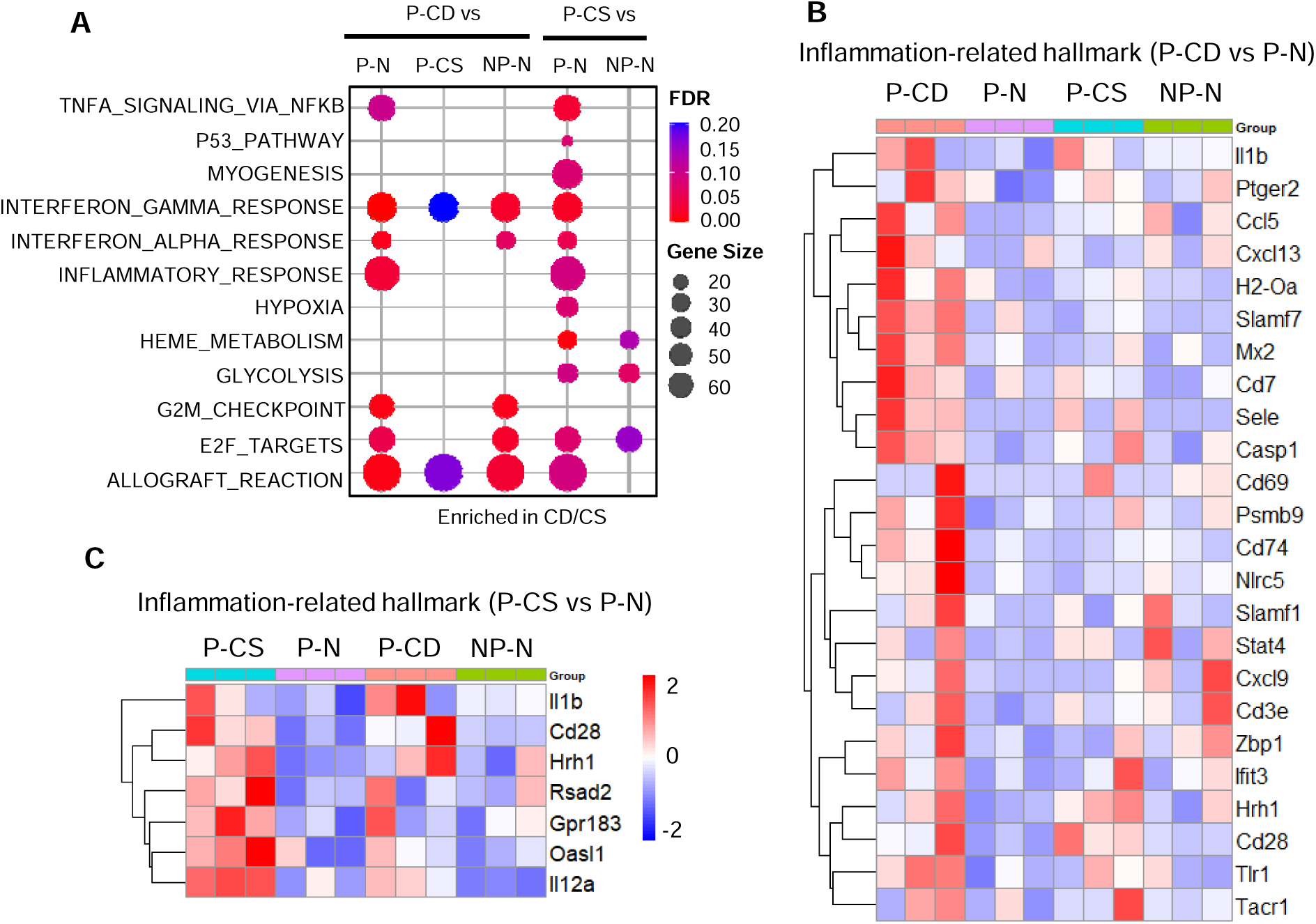
E**n**richment **of pro-inflammatory gene sets in choline-deficient hippocampus (A)** Enrich hallmarks (p < 0.05, FDR < 0.25) in P-CD and P-CS mice compared to all groups. The comparison between P-CS and P-N mice shows more enrichment gene sets, followed by P-CD versus P-N mice. **(B,C)** The heatmap of the combined immune related gene sets (TNFA_SIGNALING_VIA_NFKB, INTERFERON_GAMMA/ALPHA_RESPONSE, INFLAMMATORY_RESPONSE, and ALLOGRAFT_REACTION (p < 0.05) enriched in P-CD mice compared to P-N mice **(B)** (n = 24) and enriched in P-CS compared to P-N mice **(C)** (n = 7), respectively.

## Discussion

Women commonly experience cognitive changes during pregnancy, such as forgetfulness. The present study evaluated whether the availability of nutrients during pregnancy influences cognitive function in pregnancy and in old age. The study revealed three salient findings. First, the plasma levels of many nutrients including amino acids, glucose, lipid, and particularly choline and its derivatives were reduced during pregnancy in mice on a standard laboratory diet. This suggests that the foetal demand for nutrients imposes a strain on maternal nutrient supplies. Second, pregnant mice had impaired cognition compared to non-pregnant mice, which is worsened by choline deprivation and alleviated by choline supplementation. It is plausible that decreased choline contributes to the reported memory deficits in pregnant women. Third, multiparity together with choline availability during pregnancy have a lasting effect on the brain in old age. Multiparous mice given a choline supplement appeared healthier and performed modestly better in the MWM than those given normal or choline-deficient diets, with this trend particularly evident during the probe session among the pregnant mice, although it was not statistically significant. The RNA-Seq analysis of the hippocampus showed distinct gene signatures across the groups with different diets. The hippocampi of mice with choline deprivation showed more upregulation of genes related to immune response, whereas the hippocampi of the mice with choline supplement showed upregulated of genes including *Prl*, *Gh*, *Hba*, and *Hbb* that are positively related to spatial learning and brain health [71, 90–96]. Together, these findings show that choline supplements during pregnancy benefit women’s cognitive health both during pregnancy and later in life.

Pregnancy induces significant physiological and metabolic changes to meet the nutritional demands of the developing foetus and mother. Plasma concentrations of several essential nutrients such as iron, folate, vitamin B12, calcium, vitamin D decrease during pregnancy. Hence, dietary supplement of these key micronutrients is widely recommended [97, 98]. However, a systematic study of changes in plasma metabolites during pregnancy in humans or animal species has been lacking. A recent study reported genetic associations of plasma metabolites in early pregnancy, which offers a glimpse into the large variations among individuals and the genetic influences [99]. But this study did not investigate the differences between pregnant and non-pregnant states. Mouse model of metabolic studies has distinct advantages such as similar genetic background, controlled diet and entrained circadian rhythms. Our study obtained highly conclusive data that showed broad decreases in the levels of plasma metabolites during pregnancy, particularly choline and its related metabolites. It would be interesting to see if similar changes occur in humans, in which diet, genetic backgrounds and socioeconomic status influence metabolites changes.

It is known that choline deficiency can negatively impact brain health, particularly memory and cognition [100, 101]. Our study provides evidence to show the impact of choline availability during pregnancy on the maternal cognitive health during pregnancy and in old age. Although the observed differences in some behavioural tests were relatively moderate and, in certain cases, only marginally significant, these findings are not unexpected given that the mouse model did not have any predisposition to neurodegenerative diseases, and dietary choline deprivation or supplementation was only short term (during pregnancy). In the case of choline deprivation, mice were given a normal diet on GD5 and 9 of pregnancy to provide relief, which might explain the partial rather than robust statistical significance in some measures. Nonetheless, the MWM still demonstrated distinct differences between choline deficiency and supplementation during pregnancy, with effects continuing through to 15 months of age, although to a lesser extent. Furthermore, choline availability significantly altered gene signatures in the hippocampus in old age, confirming the lifelong impact of choline availability during pregnancy on the hippocampus.

Our findings are especially relevant in relation to the aetiology for higher risks of AD in women. There is growing evidence that choline deficiency contributes to AD pathogenesis by disrupting neuronal membrane integrity, neurotransmitter synthesis, and epigenetic regulation. Low plasma levels of choline were correlated with increased accumulation of amyloid plaques and tau tangles in people with AD [102]. Mouse studies have also shown that a choline deficient diet aggravates AD-related pathologies including elevating the level of amyloid-beta protein and altering tau protein [102]. Accordingly, choline consumption has been positively correlated with cognitive performance and risk of dementia among elderly [103, 104]. Furthermore, lifelong choline supplement ameliorated AD pathology and improved cognition in an APP/PS1 mouse model of AD [105]. It is plausible that inadequate choline intake during pregnancies also poses increased risks for dementia at the old age. Choline supplements during pregnancy could provide important cognitive benefits for both the offspring and mother.

## Materials and Methods

### Subjects

Twelve-week-old C57BL/6J mice were housed in individually ventilated cages with corncob bedding under a 12-hour light/dark cycle (lights off at 7:00 pm). All experimental procedures were conducted in accordance with the appropriate guidelines and regulations. The study protocol was approved by the Institutional Animal Care and Use Committee (NTU-IACUC), Nanyang Technological University (NTU), Singapore (IACUC Protocol Number: A22052).

### Experimental design

Mice were assigned to four different groups: non-pregnant control with a normal diet (NP-N), pregnant control with a normal diet (P-N), pregnant mice with a choline-supplemented (6g/kg choline) diet (P-CS), and pregnant mice with an intermittent choline-deprived (0g/kg) diet (P- CD). The choline diet was customized from the normal maintenance diet from Altromin (catalogue no. #1324), which is the standard diet for mice used in the animal facility at NTU. The choline diet was provided only during the pregnancy period from GD2 to 18. Due to choline deficiency having adverse effects on mouse skin, we implemented a regimen of two days on and one day off for the choline-deficient diet until GD12 (normal diet was given on GD1, 5, and 9), followed by a full choline-deficient diet until GD18. In total, there were four pregnancies throughout the experimental period. Behavioral studies were conducted during the first and fourth pregnancies, as well as at 12 and 15 months of age. The behavioral tests included open field test (OFT) for locomotor and exploratory behavior assessment, novel object recognition (NOR) and Morris water maze (MWM) for cognitive function during pregnancy and old age, and light-dark test (LDT) and forced swim test (FST) to assess anxiety and behavioral despair during the postpartum period after the fourth pregnancy.

### Behavioural testing and evaluation

All behavioral tests were conducted during the daytime in a quiet room with yellow lighting. Mice were acclimated to the room for at least 30 minutes before testing. The arena was cleaned with 70% ethanol and dried before each new subject to prevent any carry-over odors that could affect the results. All tests were recorded using ANY-maze software, except for the Morris water maze, which was recorded using EthoVision XT.

### Open field test (OFT)

The test was conducted during the first and fourth pregnancies, and at 12 months of age. A 45 cm x 45 cm open field box was used as the arena for the OFT test. Mice were gently placed in the central area of the open field box and allowed to explore the field freely for 10 minutes. The distance traveled and the time spent in both the outer and inner zones were recorded and analyzed to assess locomotor function and anxiety levels between groups.

### Novel object recognition (NOR)

The same arena used in the OFT was employed for the NOR test to ensure the mice were familiar with the environment. The procedure consisted of two phases: a training trial followed by a testing phase. During the training trial, two identical objects were placed in opposite corners of the arena and mice were allowed to explore for 3 minutes. After one hour, one of the identical objects was replaced by a novel object, and mice were returned to the arena and allowed to explore for another 3 minutes. The novel object had a similar size to the previous object, but with a different color and shape. To prevent movement, the objects were affixed to the arena floor. To eliminate any potential olfactory cues, the objects were wiped with 70% ethanol, followed by water, and then dried before being used with the next mouse. The exploration time spent on the familiar and novel objects was recorded and analyzed. The discrimination index was calculated using the following formula:

Discrimination index = (Novel object time spent − Familiar object time spent) / (Novel + Familiar object time spent)

### Morris water maze (MWM)

A circular pool with a diameter of 1.2 m, filled with water at 22°C–23°C, was used for the MWM test. To provide better colour contrast with the black C57BL/6J mice, non-toxic white paint was used to color the water. The pool was divided into four quadrants in the software, arbitrarily labeled as north (N), south (S), east (E), and west (W). A hidden platform was positioned at the center of the SW quadrant, approximately 1 cm below the water surface. The MWM test consisted of two phases: four days of training (with the hidden platform) followed by a probe session (no platform) on the fifth day. Four different sets of curtains were hung around the pool to serve as visual cues to help the mice remember the platform’s location. To reduce anxiety during their first exposure to water, pre-training was conducted by allowing the mice to explore without data recording.

During the training phase, each mouse was trained four times a day. The mouse was gently placed into the pool facing the wall at one of the following starting positions: N, E, NW, or SE. The mice were allowed a maximum of 60 seconds to find the hidden platform. If the mouse successfully found and stayed on the platform for 10 seconds, the software was stopped and the escape latency recorded. If the mouse failed to find the platform after 60 seconds, it was guided to the platform. Whether successful or not, the mouse was allowed to remain on the platform for 60 seconds to learn and remember its location.

During the probe session, the hidden platform was removed and mice were placed at the NE position and allowed to swim in the pool for 60 seconds. The software recorded the time it took for the mice to first enter the platform’s location.

### Light-Dark test (LDT)

The light-dark box consists of two compartments: an exposed light compartment with white walls and a dark compartment with black walls, enclosed on all sides except for a small hole that allows the mouse to move freely between the compartments. The mouse was placed in the center of the light compartment and allowed to move freely within the box for 5 minutes. The time spent in both the light and dark zones was recorded to compare anxiety levels between groups.

### Forced swim test (FST)

A transparent Plexiglas cylinder (30 cm height x 10 cm diameter) filled with water at 22°C– 23°C was used to conduct the forced swim test. The mouse was placed in the water for 6 minutes. Both active (swimming) and immobile (little or no movement, solely to keep the nose above the water) states were recorded. The water was changed before the next mouse.

### Plasma collection and NMR Analysis

Mice on gestation day 1 (GD1) and GD7.5 were anesthetized, and blood was collected via cardiac puncture in heparinized tubes. The collected blood was centrifuged at 10,000 rpm, 4°C for 10 minutes. The plasma was transferred to a new Eppendorf tube and snap-frozen in liquid nitrogen. The non-targeted NMR analysis was carried out at the Singapore Phenome Centre (SPC)of Nanyang Technological University. A total of 50 μL of serum sample were added with 30 μL water followed by addition of 75 μL of PBS D2O buffer (0.15 M KH2PO4 phosphate salt) and 3-Trimethylsilyl propionic acid-d4 (TSP) sodium salt (pH=7.4). The sample was vortexed and followed with centrifugation for 10 mins before 150 μL sample was transferred to 3 mm NMR tube. CPMG (Carr-Purcell-Meiboom-Gill) spectra were acquired from 600 MHz NMR spectrometer (Ascend III HD, Bruker), equipped with 5mm BBI 600 MHz Z- Gradient high-resolution probe (Standard Probe) at 310 K. 90o pulse was set ∼10 μs and total of 120 scans were accumulated into 20 ppm spectral width. Water peak pre-saturation and a spin-spin relaxation delay were set to 100 ms. Metabolites were extracted from spectra and the concentration was calculated based on the TSP peak.

### RNA-Seq analysis of gene expression

RNA-seq analysis was performed to determine the effect of pregnancy and choline diet on global gene expression in mouse hippocampus. The hippocampus was harvested from mice at 15 months of age and snap-frozen in liquid nitrogen. The hippocampus was homogenized and total RNA was extracted using TRIzol Reagent (Thermo Fisher Scientific, USA), followed by DNase I treatment using a DNA-free™ DNA Removal Kit (Invitrogen, Carlsbad, CA, USA). Total RNA samples were sent to the Genome Institute of Singapore, Agency for Science, Technology, and Research, for library preparation and paired-end sequencing.

The FASTQ files were trimmed using Trim Galore, followed by quality control with FASTQC. STAR was used to map the reads and FeatureCounts was used to generate gene counts. Gene counts were subjected to DESeq2 analysis and normalized counts were used for subsequent analyses.

### Data Availability

The RNA-seq data that support the findings in this study is available in NCBI Gene Expression Omnibus (GEO) under accession number GSE293570 (https://www.ncbi.nlm.nih.gov/geo/query/acc.cgi?acc=GSE293570).

### Data filtering of RNA-seq results

A total of 10 comparisons were made between groups. The heatmap was generated using the pheatmap package in R to display the significantly differentially expressed genes. For the heatmaps in the RNA-seq analysis overview, significant gene lists with a p-value < 0.01were selected. Genes were further filtered based on the following criteria: (1) inconsistent expression among the three biological replicates based on log2 TPM values, and (2) one group having zero counts for all replicates, while the other group has an average count of fewer than 10.

### Gene set enrichment analysis (GSEA)

The official software was downloaded from https://www.gsea-msigdb.org/gsea/index.jsp, and used to perform GSEA analysis using the default settings. A hallmark was considered significantly enriched for a p-value < 0.05 and FDR < 0.25. A bubble plot of the GSEA results was created using the ggplot2 package in R. Genes enriched (p < 0.05) in the selected hallmark were extracted and displayed in the heatmaps.

### Real-time PCR analysis

2μg of DNAase-treated RNA from each hippocampus was reverse transcribed into cDNA using Superscript II reverse transcriptase (Invitrogen, Carlsbad, USA). Real-time PCR was performed with KAPA SYBR® FAST qPCR Master Mix (Kapa Biosystems, Wilmington, USA) on QuantStudio™ 6 Flex Real Time PCR System (Applied Biosystems, Foster City, CA). The real-time PCR primers used were: Growth hormone (GH) forward 5′- AAGAGTTCGAGCGTGCCTAC-3′ and reverse 5′-GGATGGTCTCTGAGAAGCAGA-3′, Prolactin (PRL) forward 5′-ATCAATGACTGCCCCACTTC-3′ and reverse 5′- CTGCACCAAACTGAGGATCA-3′, Hemoglobin Subunit Alpha (HBA) forward 5′- TCCCGTCAACTTCAAGCTC-3′ and reverse 5′-GTGCTCACAGAGGCAAGGA-3′, Hemoglobin Subunit Beta (HBB) forward 5′-AGGTGAACGCCGATGAAGTT-3′ and reverse 5′-ACTTTCTTGCCATGGGCCTT-3′, 36B4 forward 5′- GATCGGGTACCCAACTGTTGCC-3′ and reverse 5′- CAGGGGCAGCAGCCGCAAATGC-3’ . The delta Ct method was used to calculate the relative expression levels of the four interested genes by normalizing against the 36B4 Ct values.

### Statistical analysis

All statistical analyses were conducted using GraphPad Prism 9.5.0. Data presented in the figures are shown as mean ± S.E.M. Analysis has been done based on two categories: non-pregnant (NP-N) versus pregnant mice (P-N) and between all pregnant groups (P-N versus P- CS versus P-CD). Parametric tests were applied as most of the behavioral test data was normally distributed. Two-way ANOVA was used when accounting for two variables in a particular parameter (e.g., variable 1: outer and inner zones in the open field box, variable 2: four mice groups, parameter: time spent in the zones). Repeated measure was applied when the mice were subjected to the same test across different days in Morris Water Maze. One-way ANOVA was used when comparing only one variable across more than two groups.

Student t-test was used when comparing two groups, within groups, or between two zones or objects. For the NMR results, one-way ANOVA was used to analyze the differences between the following groups: WT non-pregnant, WT pregnant, PgR mutant non-pregnant, and PgR mutant pregnant. The qPCR gene expression results were analyzed by one-way ANOVA between groups. However, data for GH and PRL were not normally distributed, hence, Kruskal-Wallis test was used to analyze these two genes. Tukey’s or Sidak post-hoc tests (Dunn’s test was used for non-parametric data) were performed if any significant differences were found in the analyses. All p-values < 0.05 were considered statistically significant. Any outliers identified through Grubbs’ test were removed from the analyses.

