## Supplementary Figures for "Impact of Choline intake during pregnancy on maternal cognition and hippocampal gene signature in old age"

**Supplementary Fig. 1 Pregnancy is associated with significant decreases in the plasma levels of metabolites, particularly choline-related metabolites**

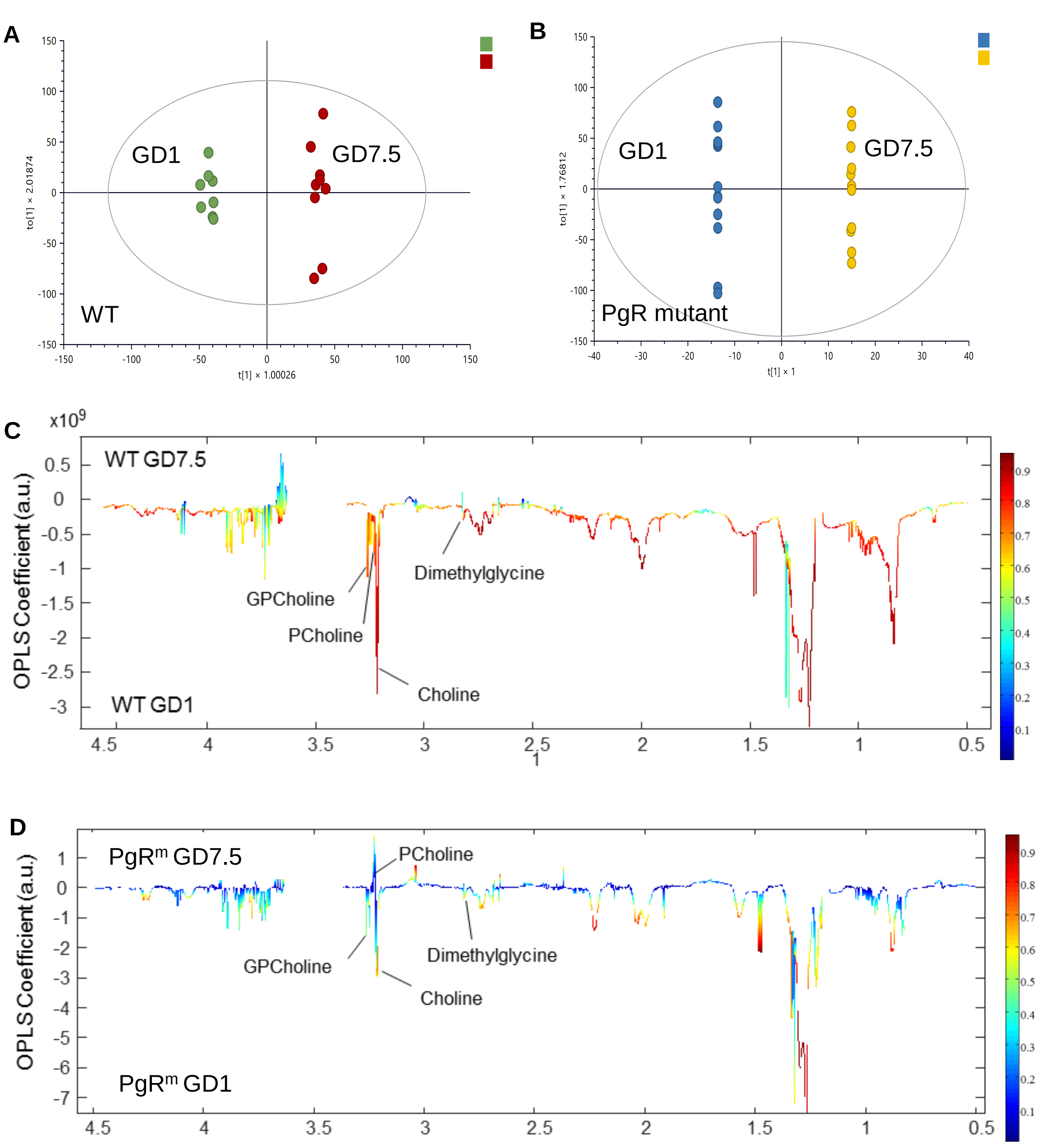


**Supplementary Fig. 1 Pregnancy is associated with significant decreases in the plasma levels of metabolites, particularly choline-related metabolites (A)** The OPSL-DA score plot clearly shows the data points are separated into two clusters in the wild type (WT) mice, indicating significant changes in plasma levels of metabolites between GD1 and GD7.5. **(B)** The OPSL-DA score plot shows much lower degree of separation on the X axis, indicating less metabolic changes between GD1 and GD7.5 in PgR mutant mice. **(C,D)** OPLS-DA coefficient plots generated from NMR spectra of serum samples showing changes in the plasma levels of choline and its derivatives during early pregnancy. Peaks pointing upward indicate increased metabolites in GD7.5 or in dpc7.5 compared with GD1 or dpc1, respectively, whereas peaks pointing downward indicate decreases. The color of the spectra indicates the significance of the changes with a warmer color having more significance than a colder color, as indicated in the colored bar. There was significant reductions in the levels of choline, phosphatidyl choline (PC), glycerylphosphorylcholine (GPC), and dimethylglycine at GD7.5 compared to GD1 in WT mice **(C)**. In contrast, these metabolites were not significantly changed in PgR mutant mice (**D)**.

### Supplementary fig. 2 Mice tend to show progressively slower swimming speeds over the training days in the Morris Water Maze

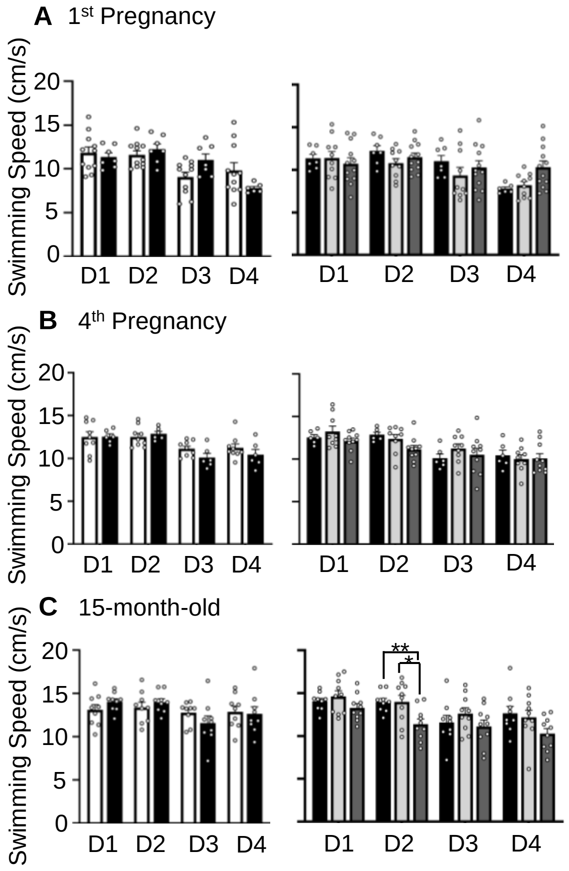


**Supplementary Fig. 2 Mice tend to show progressively slower swimming speeds over the training days in the Morris Water Maze (A,B,C)** Bar charts showed swimming speed between NP-N and P-N mice, and between pregnant groups on each training days during first pregnancy **(A)**, fourth pregnancy **(B)** and at 15 months old **(C)**. Although the P-CD mice showed significantly slower swimming speed on training day 2 at 15 months old, this was not accompanied by a longer escape latency to find the hidden platform on the same day. Mean ± SEM is used to present all numerical data. The degree of statistical significance is indicated with asterisks (* $p < 0.05$ ,** $p < 0.01$).

**Supplementary Fig. 3 All mice groups exhibited same degree of depression in Forced Swim Test**

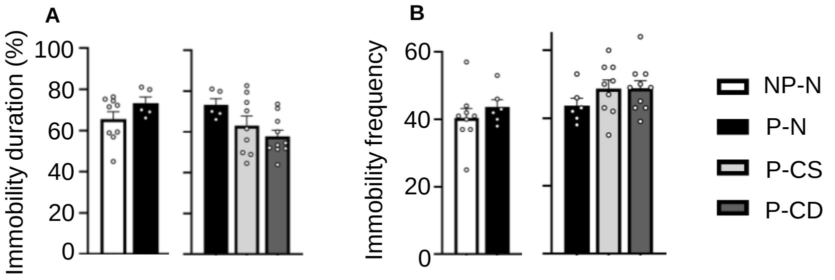


**Supplementary Fig. 3 All mice groups exhibited same degree of depression in Forced Swim Test (A)** The percentage of immobility duration during the forced swim test did not show any differences between groups. The depression levels are similar across groups. **(B)** The frequency of immobility during the forced swim test also showed no differences between groups. Data are presented as mean ± SEM.

### Supplementary fig. 4 Choline-deficient diet during pregnancy causes hair loss during pregnancy and in old age

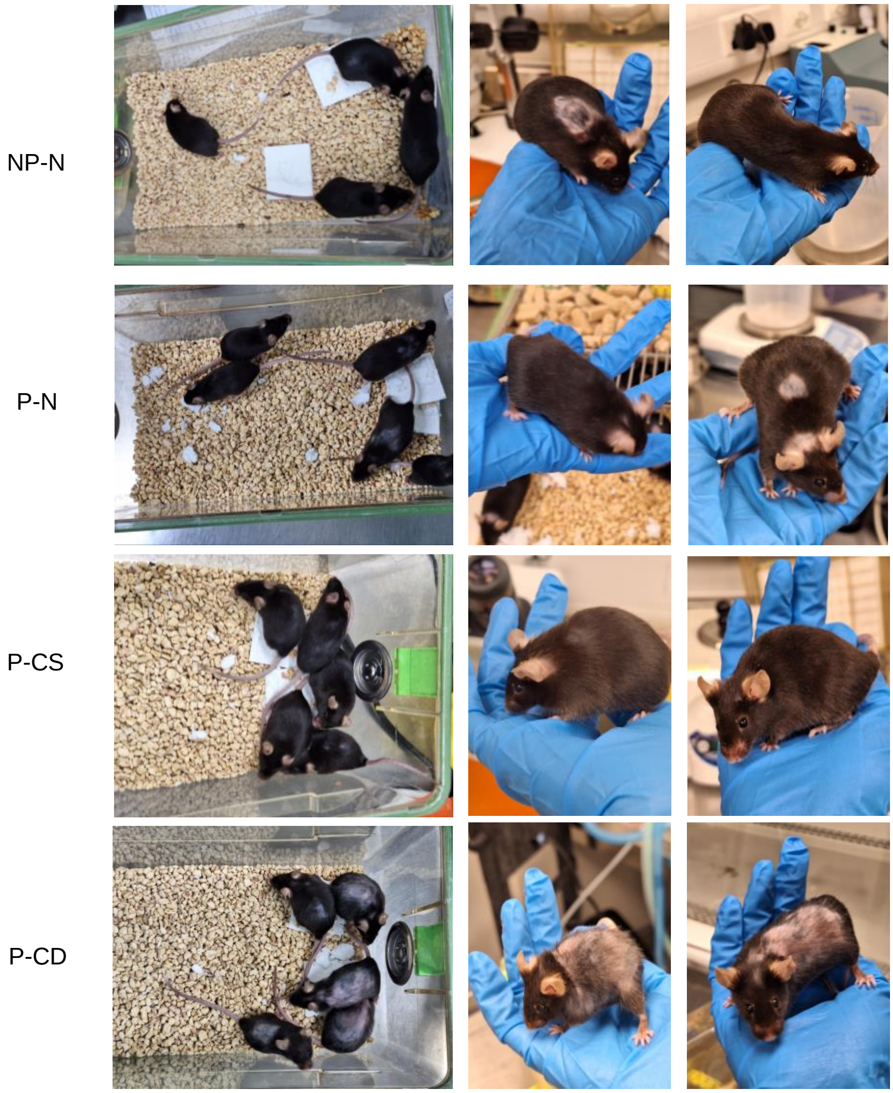


**Supplementary Fig. 4 Choline-deficient diet during pregnancy causes hair loss during pregnancy and in old age**
The pictures show the appearance of mice at 12 months of age. P-CD mice had a noticeable alopecia condition. Note that ‘barbering behavior’ (in which the dominant mouse will tend to nibble on the fur of the other mice repetitively in the same area of skin, and leading to patchy fur loss appearance) was observed in all groups.
